# Linking acoustic telemetry data to spatial covariates in river networks with spatially explicit capture-recapture models

**DOI:** 10.1101/2025.02.09.637304

**Authors:** Joseph R. Bottoms, Marie Auger-Méthé, Bryce O’Connor, Michael Power, David A. Patterson, J. Mark Shrimpton, Steven J. Cooke, Eduardo G. Martins

## Abstract

1. Spatially explicit capture-recapture (SECR) models extend classical capture-recapture models to include spatially-explicit animal locations and environmental covariates. SECR models have been widely employed in terrestrial studies to predict the population size and densities of animals assuming a closed population over a defined area. In this work, we extend and apply SECR models to a novel use-case that both uses a relative density formulation and accounts for biased tagging distributions to estimate parameters of habitat use from acoustic telemetry data in a branching river network.
2. Using SECR models, we predict how temperature distributions during the summer feeding season influenced how tagged Arctic grayling (*Thymallus arcticus*) distributed themselves through the Parsnip watershed in northcentral British Columbia, Canada. We found that the relative density of tagged Arctic grayling peaked at water temperatures of 12.4 °C. In warm years, relative densities were constricted as parts of the watershed became unfavorably warm. In cool years, fish were distributed widely throughout the watershed.
3. In acoustic telemetry, only the tagged population is available for detection. We highlight several specific considerations and assumptions for using this approach: i.e. (a) activity centres are assumed to remain in the same location throughout the study period, thus the time window of the study should be selected accordingly (e.g., exclude migratory periods); (b) inferences from acoustic telemetry data depict relative (not absolute) densities; and (c) spatial tagging effort must be defined in the model to ensure that predictions are not merely an artefact of tagging effort across space and time. When applied following these assumptions, this method is broadly useful for ecologists as it presents a quantitative way to merge automated telemetry datasets with discrete habitat parameters that drive population distributions through time and are relevant to managers and conservation professionals. Further, this method can be applied to branching river networks in which topological challenges have hindered other statistical approaches.

## Introduction

Spatially explicit capture-recapture (SECR; alternately Spatial Capture-Recapture or SCR) is a class of hierarchical statistical models which extends classical capture-recapture models to include both georeferenced animal locations and covariates (Borchers and Efford 2008, Efford and Fewster 2013, Royle et al. 2014). Compared to their predecessors that do not accommodate the spatial structure of the data, SECR models are used by ecologists to estimate the population size and density of a population of animals ranging freely around a fixed trapping array (Borchers & Efford 2008; Royle et al. 2014). These versatile models can accommodate observation data from any source in which individuals can be uniquely and repeatably identified at discrete locations in space, including (but not limited to) camera traps, acoustic surveys, and hair snares (Royle et al. 2014). SECR models have been applied most frequently to data collected from terrestrial organisms moving freely in a two-dimension (2-D) state space (i.e. the collection of pixels representing all possible locations for activity centres across a geographic area; Royle et al. 2014).

Fewer studies have applied SECR to aquatic systems. Those that have done so have typically modeled movements along one functional dimension (1-D; upstream and downstream along a single river channel; Raabe et al. 2013; Haydt et al. 2022), with limited extensions into the 2-D aquatic environment (Marques et al. 2012; Pirotta et al. 2014). Applications of SECR models to data collected in river networks are even less common (a limited network was used in Murphy et al. 2021) because a branching (dendritic) system cannot be easily represented by one or two dimensions. While a state space represented in 1-D cannot accommodate branches in the river system, a 2D state space which captures the extent of a watershed will be dominated by pixels of habitat which are non-traversable for strictly aquatic species (i.e., land).

In terrestrial applications, modeling how animals move in a dendritic network has been addressed using ecological distance models (Sutherland et al. 2015). However, this solution is unnecessarily complex for strictly aquatic species. The ecological distance model allows for terrestrial organisms to limit movement across undesirable habitats but does not completely restrict it, whereas strictly aquatic organisms simply cannot be on land. A more suitable approach to modeling the activity centres of aquatic animals moving within a dendritic state space is offered in the `secrlinear` package for R statistical software (Efford 2023; R Core Team 2023). The package, which is a wrapper for the 2D parent package `secr` (Efford 2023), applies a linear mask approach in which the dendritic network is reduced to a series of 1D lines connected at confluence points within a 2D space, effectively eliminating all land pixels from consideration. From this, a pairwise distance matrix between all wetted pixels substitutes the Euclidian distance generally used in these models (Efford 2023).

Acoustic telemetry (hereafter AT) has been largely employed in aquatic systems and forms the backbone of many large-scale collaborative monitoring programs in both the freshwater and marine environments (e.g., the Ocean Tracking Network, European Tracking Network, the Great Lakes Acoustic Telemetry Observation System, and the Integrated Marine Observing System, among others; Hussey et al. 2015; Hoenner et al. 2018; Hostetter and Royle 2020; Alós et al. 2022). AT datasets are generated by tagging individuals with an acoustic transmitter and releasing them among an array of acoustic receivers. The receiver array then monitors the approximate spatiotemporal locations of each tagged individual by receiving, decoding, and recording signals from transmitters that pass within the detection range of the receivers (Whoriskey et al. 2019). While AT enables monitoring of aquatic organisms at spatial scales that would not be feasible using other sampling methods, it cannot detect new, untagged individuals within the vicinity of the receivers (as opposed, for example, to camera traps used to study individuals with uniquely identifiable markings, e.g. Dorazio and Karanth 2017). Because the tagged population may not be representative of the untagged population, SECR models fit to AT data alone cannot estimate population size, though they can be suitable for modeling the relative density of a tagged population in relation to spatial covariates.

This paper applies SECR models modified for use with AT data collected in a dendritic river network to quantify the relative density of tagged individuals in relation to spatial covariates (which is conceptually analogous to second-order habitat selection; *sensu* Johnson 1980). Through a case study, we demonstrate how quantifying these relationships with spatial covariates can be used to explore ecological hypotheses. Specifically, using the packages `secr` and `secrlinear` (Efford 2023), we assess whether the seasonal distributions of Arctic grayling (*Thymallus arcticus,* Pallus 1776) in the Parsnip River watershed of north-central British Columbia, Canada, are influenced by watershed-scale temperatures. As high temperatures likely limit the abundance and distribution of summer rearing Arctic grayling (Stamford et al. 2017), we expected to find a negative response between high temperatures and the activity centres of tagged Arctic grayling. We conducted our analysis across an ecologically-relevant period when Arctic grayling use spatially discrete summer rearing habitats (Blackman et al. 2002; Stamford et al. 2017).

## Materials and Methods

### SECR models and their application to AT in dendritic networks

The data collected by AT are time-stamped detections of *n* individuals at *K* receivers. For convenience, we consider daily time intervals and discard repeat detections within each day. The data are then binary values *y*_*ijk*_ indicating whether the tagged individual *i* was detected during occasion (day) *j* at acoustic receiver *k*. We use ***yi*** for the matrix of data for individual *i* and ***y*** for the combined dataset. We consider each fish to occupy a contiguous and stationary home range within a dendritic river system; the location of a home range is indicated by its activity centre. Our goal is to describe the distribution of activity centres within the river system as a function of spatial covariates (second-order habitat selection in the terminology of Johnson 1980). This is equivalent to modelling the density of activity centres using, for example, a log-linear model

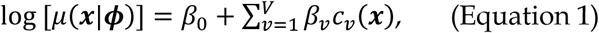

where *μ*(***x***) is the density (e.g. the expected number of activity centres per km) at location ***x***, *c*_*v*_(***x***) is one of *V* covariates measured at ***x***, *β*_0_ is the intercept, and *β*_*v*_ are coefficients to be estimated (denoted together with *ϕ*).

Eq. 1 cannot be fitted directly to AT data for multiple reasons, some of which are specific to AT and riverine systems. First, activity centres are latent and not observed, an issue that can be easily addressed with spatially explicit capture–recapture models (SECR; Borchers and Efford 2008, Efford and Fewster 2013, Royle et al. 2014). Latent activity centres are modelled in SECR by integrating the likelihood over all potential locations. Second, the Euclidean distances commonly used in SECR are not appropriate for a dendritic network. We replace them with run-of-the-river distances computed using standard algorithms for discretized graphical networks (Csardi and Nepusz 2006). Third, unlike camera traps or hair snares that can detect previously unmarked individuals, acoustic receivers can only detect tagged individuals. This limitation of AT has two important impacts on the SECR model. Tagged fish may differ in their spatial distribution from the population at large because capture for tagging can be spatially selective (e.g., dictated by access to capture sites and the fraction of the population that was tagged cannot be determined, therefore the absolute density (or total population) cannot be estimated. We use a newly developed SECR model that handle these issues by including a distance-related decline in probability of detection that is linked to capture effort and focuses on relative density, described by the coefficients *β*_*v*_, and ignores the intercept in Eq. 1 (Efford 2025).

Consider a hypothetical one-phase SECR study (i.e., a study in which all fish could be detected without a separate tagging process) and *n* fish are detected at least once. Parameters of both detection (*θ*) and the distribution of fish activity centres in the river system (*ϕ*) can be estimated by maximising the likelihood

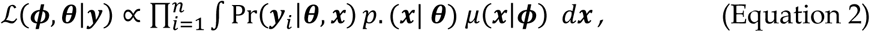

where Pr(***y***_*i*_|*θ*, ***x***) is the probability of observing the detection history of fish *i* if its activity centre is at ***x***, *μ*(***x***|*ϕ*) is the density of activity centers at ***x*** (Eq. 1), and *p*. (***x***|*θ*) is the probability that a fish with an activity centre at ***x*** will appear in the dataset. This last term can be viewed as a spatial filter that can allow the activity centre distribution of the detected fish to differ from the population as a whole. The integration is over the extent of potential habitat (activity centre locations), and observations ***y*** provide information on the detection process (parameter vector *θ*) that is shared between components of the likelihood. (Efford 2025).

In a study in which there are distinct tagging and acoustic detection phases (i.e., a two-phase study), it is inappropriate to estimate *p*. (***x***) from AT data alone because in such a study it represents the unobserved probability that a fish with an activity center at ***x*** was captured and tagged. However, one can use external information (e.g., tagging effort) to create an estimate of *p*. (***x***) that we denote with *P*_*T*_(***x***) and use it as a substitute in Eq. 2 (Efford 2025).

In our case study, we fitted a half-normal detection function to the AT data (though the use of other functions is possible), using the run-of-the-river distance *dik* between the hypothetical activity centre of fish *i* and the location of receiver *k*: Pr(*y*_*ijk*_ = 1|*θ*, ***x***) = *g*_0_ exp[−*d*_*ik*_^2^ /(2*σ*^2^)], where *θ* = (***g*_0_**, *σ*) were parameters to be estimated. The run-of-the-river distance *dik* here is the critical term for applying SECR models to dendritic river networks.

### Study Area

Our case study data were collected in the Parsnip River watershed (approximately 5,000 km^2^) in north-central British Columbia, Canada (Figure 1). The area is in the overlapping traditional territories of Treaty 8 signatory Nations (Prophet River, Saulteau, and West Moberly First Nations) and the traditional and unceded territory of the McLeod Lake Tse’Khene. The Parsnip River is a moderate- to low-gradient system that flows northwest along the Rocky Mountain Trench. Over its 240km course, it is fed by many low-order tributaries draining the Hart Ranges as well as four higher-order rivers which support Arctic grayling and were selected for this study: the Anzac, Table, Hominka, and Missinka Rivers. Together, the Parsnip River and these main tributaries contain most of the Arctic grayling in the Parsnip River watershed (Stamford et al. 2017).

**Figure 1.**
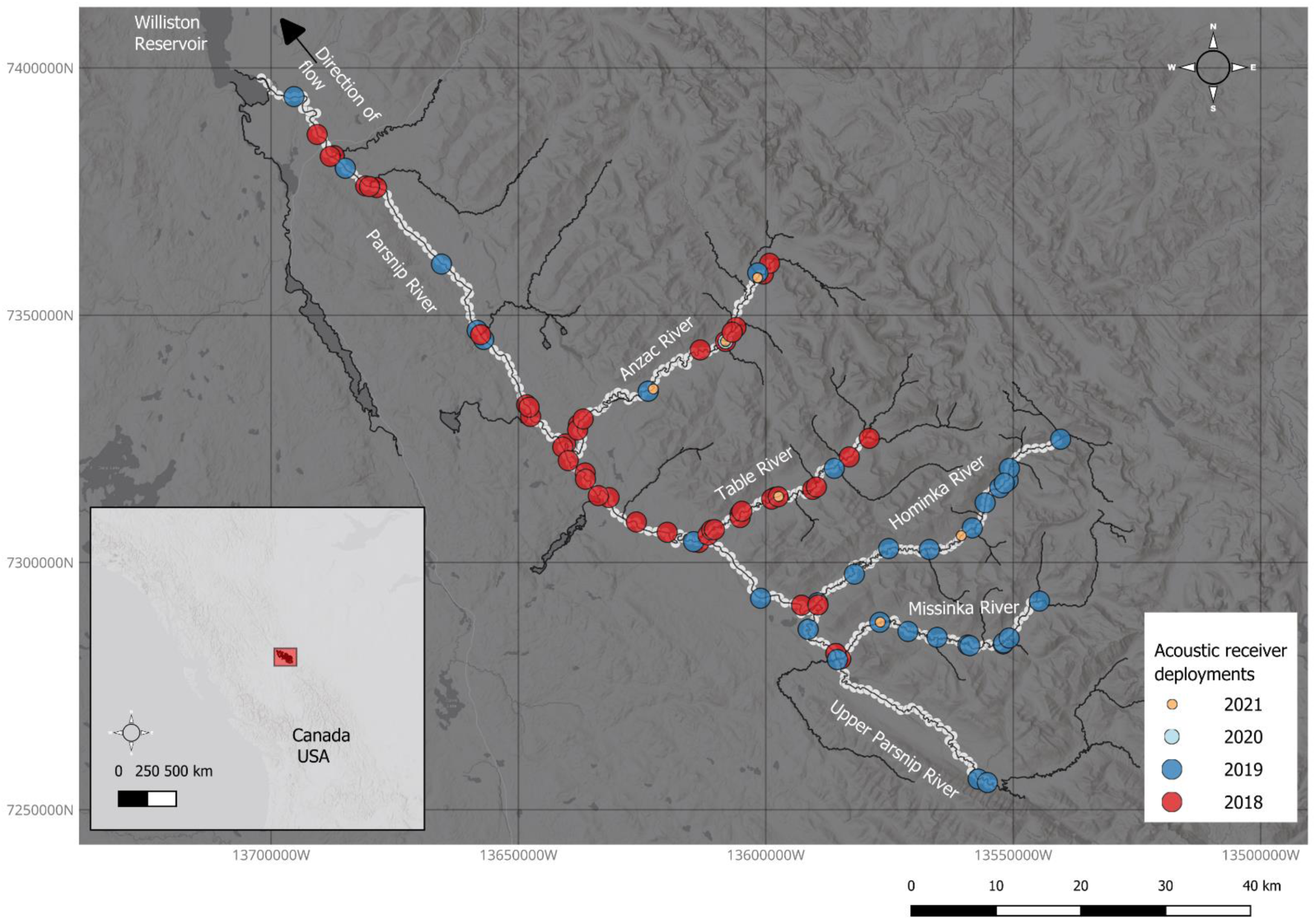
The Parsnip River watershed in north-central British Columbia, Canada, was discretized into a state-space mask of 366 pixels at a resolution of 1 Rkm/pixel. The state-space mask includes the Parsnip River mainstem and four major tributaries included in this study: the Anzac, Table, Hominka, and Missinka Rivers. The watershed flows northwest through the Rocky Mountain Trench into the Williston Reservoir. Acoustic receiver deployments are indicated by year; overlapping points indicate replacements of compromised or lost acoustic receivers.

### Acoustic tagging

Adult Arctic grayling were captured by angling with either dry flies or spin casting based on current angling conditions. Tagging began after spring freshet and continued until ice-up at various sites in the Parsnip River and its tributaries. Captured Arctic grayling > 230 g were measured for fork length (mm) and mass (g) before being surgically tagged with acoustic transmitters (Innovasea [formerly Vemco] V9 tags, 3.6 g in air, 90-150 second transmission interval, 803-859 days of battery life, Bedford, NS, Canada). All individuals were captured as authorized under permits PG18-356580, PG19-523435, PG20-606121, and PG21-622265 issued by the BC Ministry of Forests, Lands, Natural Resources Operations and Rural Development (currently BC Ministry of Water, Land and Resource Stewardship). Fish handling and tagging were conducted in accordance with the University of Northern British Columbia’s Animal Care and Use Committee (ACUC protocol 2018-06).

We initially attempted to distribute capture and tagging effort equally in the upper, middle, and lower reaches of each tributary (with some additional effort applied on the Parsnip mainstem itself). However, due to logistical constraints (e.g. road access), variable angling conditions, and heterogenous distribution of Arctic grayling along tributaries in the summer, tagging effort varied in different sections of the watershed (Table 1).

**Table 1.**
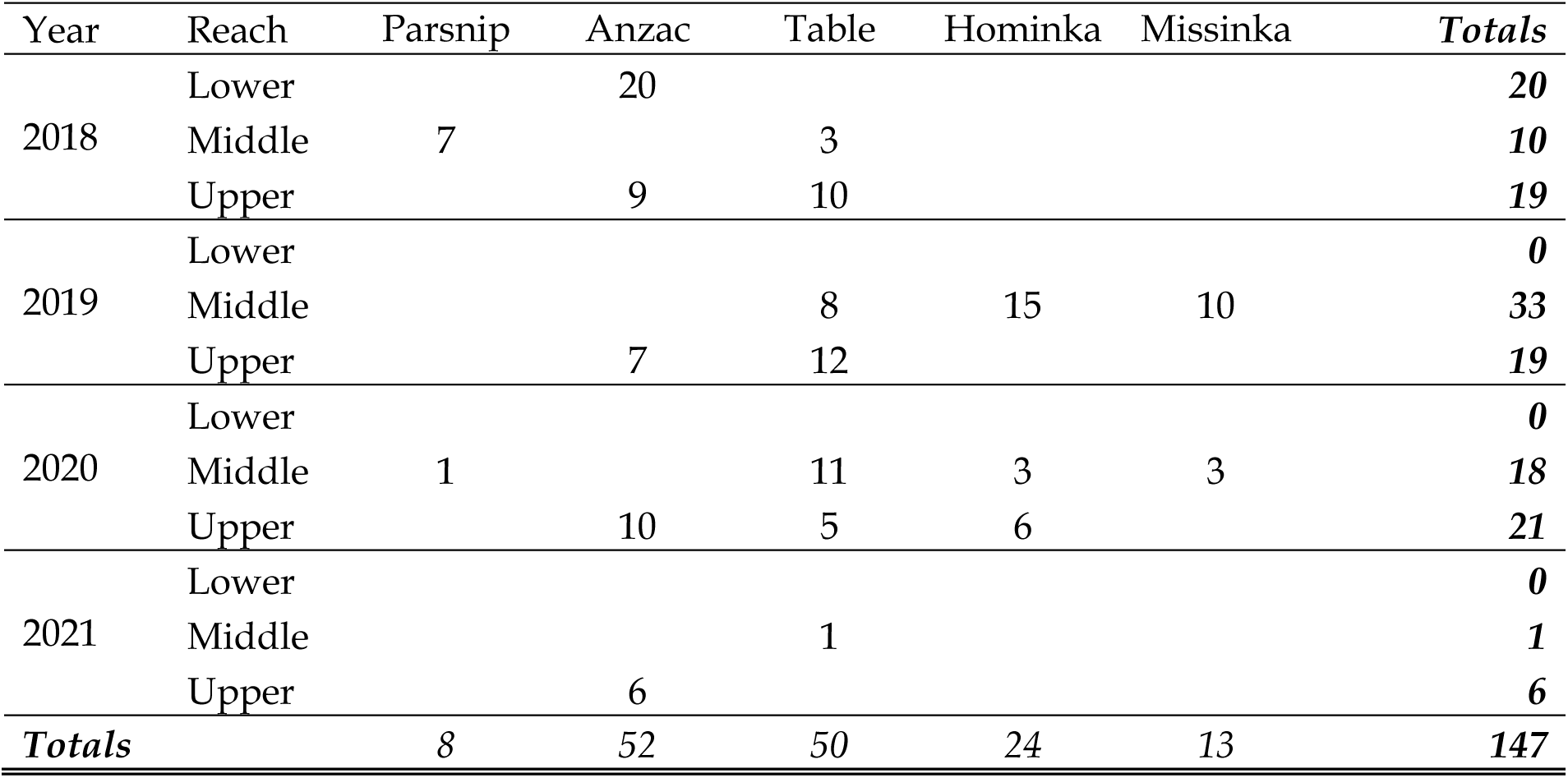
Annual counts of Arctic grayling (*Thymallus arcticus*) acoustically tagged in the Parsnip River and its four major tributaries.

### Data Collection

Innovasea VR2W- and VR2Tx-69 kHz acoustic receivers were deployed over an extent of 366 river km. The selection of deployment sites was based on site accessibility, substrate conditions (coarse enough to not lose receivers to silt compaction, while fine enough to minimize acoustic interference from water flowing over large substrates), distance from public access points (accessible but not likely to be seen by anglers), and amenability to receiver redundancy within 2 km. Receivers were deployed following a clustered strategy (2-4 receivers spaced 0.5-2 km apart), while distributing the receiver clusters widely in the study rivers to optimize data collection for use with SECR models (Royle et al. 2014). The clustered deployment also provided redundancy in the event of receiver loss or damage. The receivers were moored hydrophone-up at each site using a cement block, steel cable, and a cable anchoring system (Duckbill Earth Anchors, Fort Mill, SC).

Deployment of the acoustic receiver array began in late July 2018, with 54 receivers deployed across the watershed by the end of the open-water season. Receiver loss was prominent in the 2019 and 2020 freshets, which sustained high-water years (particularly in the Table and Anzac Rivers; *Supporting Information, Figure 5*). Deployments and replacement of lost receivers continued after freshet in 2019 (27 lost; 41 deployed; 68 total), 2020 (18 lost; 2 deployed; 52 total), and 2021 (12 lost; 10 deployed; 50 total) (Table 2). We compensated for variable receiver coverage by using seasonally specified array-deployment files indicating whether each receiver was online or offline on each occasion (see tutorial in *Supporting Information*, *Tutorial* and at <link<).

**Table 2.**
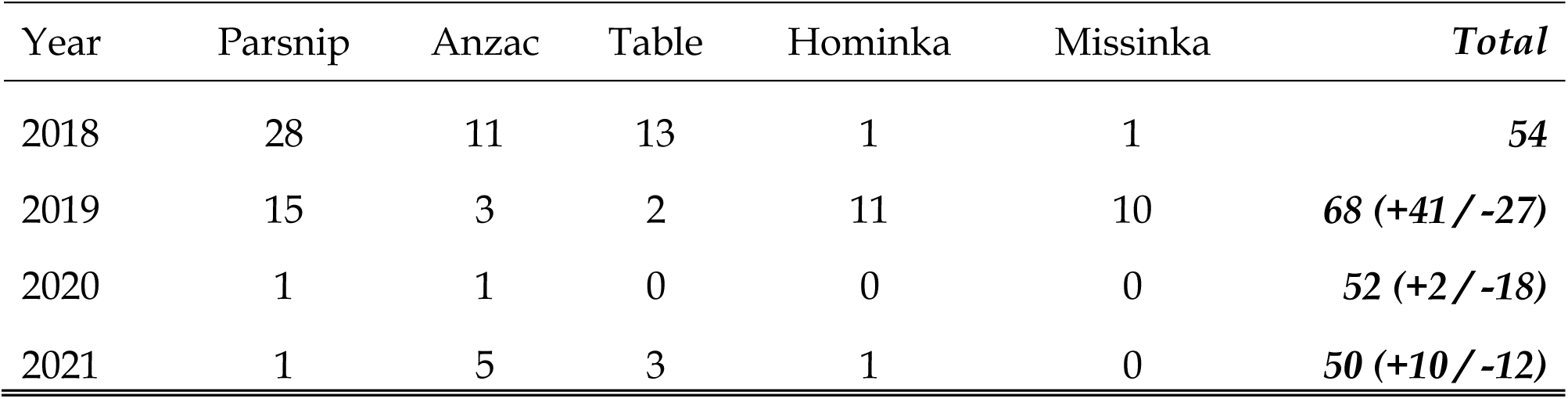
Annual deployments and active coverage extent of the acoustic receiver array by river. High receiver loss occurred during a high discharge freshet in 2020. See Figure 1 for a visual description of these data. Values in parenthesis represent the receiver deployments (+) and losses (-) during each year.

| Year | Parsnip | Anzac | Table | Hominka | Missinka | *Total* |
| --- | --- | --- | --- | --- | --- | --- |
| 2018 | 28 | 11 | 13 | 1 | 1 | **54** |
| 2019 | 15 | 3 | 2 | 11 | 10 | **68 (+41 / -27)** |
| 2020 | 1 | 1 | 0 | 0 | 0 | **52 (+2 / -18)** |
| 2021 | 1 | 5 | 3 | 1 | 0 | **50 (+10 / -12)** |

Deployed receivers were downloaded and maintained once per year during autumn base flow conditions. Equipment retrieval began in late summer of 2021 following the approximate Arctic grayling downriver migration and concluded on 30 October 2021. The data included in our model was filtered for false (erroneous tag ID) and out-of-bounds (erroneous detection date) detections with the R package `actel` (Flávio & Baktoft 2021). Skip movements (cases where a tag was detected at two receivers but missed by a receiver located between the two) were permitted to retain valid detections which would otherwise be filtered out because of variable detection efficiency. We compared the model outputs in our case study to models fit to the same dataset filtered for skip movements and both model performance and predictions were similar.

### Fitting SECR models to detection data

We used the implementation of SECR for linear habitats in the R packages `secr` 5.0.0 and secrlinear 1.1.4 (Efford 2023a,b). The river system was discretized into 1 km pixels; network topology was inferred from the proximity of pixels in Euclidean space and checked visually. Network distances were calculated by Dijkstra’s algorithm in the R package `igraph` (Csardi and Nepusz 2006).

We did not keep track of tagging effort, thus we computed *P*_*T*_(***x***), the spatial filter due to tagging, as a cumulative kernel density estimate of tagging locations in each season (minus the density of tags deployed longer than their expected battery life). The likelihood in Eq. 2 with *P*_*T*_(***x***) substituted for *p*. (***x***|*θ*) was maximised numerically in `secr` with the function `secr.fit`. This required two specialised settings: `relativeD` and `externalpdot` (see R code in *Supporting Information, Tutorial*).

SECR models can accommodate spatial covariates on the condition that they are available across the state space used to model activity centres (with additional flexibility to integrate receiver-level covariates and individual-level covariates). The models in our case study used a dendritic state-space of 366 pixels at a resolution of one river kilometer (Rkm) and covariates were defined to the same resolution. Temperature for each state-space pixel was obtained using predictions from a spatial stream network model parameterized with data collected from 221 temperature loggers installed throughout the watershed over the same spatiotemporal extent as the acoustic receiver array (O’Connor et al. *in review*). The temperature covariate represents the summer feeding period for Arctic grayling (when high temperatures have the potential to be a limiting factor to their distribution) and represents the mean pixel predictions of daily temperatures (*Temp*) from July 1 – September 15.

Arctic grayling SECR models were fit for the summer detection window (Jul. 1 to Sep. 15). As Arctic grayling seasonal distributions are well-understood and they remain close to their territorial feeding sites throughout the summer (Stamford et al. 2017), the SECR assumptions that the population is closed and that activity centre locations are fixed within the temporal windows being modelled (Efford 2023) were reasonably satisfied in our case study. We selected among three candidate models, which included the no effects model (i.e., no covariates, or the intercept-only model of activity centre density) and models including standardized temperature as both linear and polynomial predictors (*Temp* and *Temp + Temp^2^;* the polynomial term was included as there are likely upper and lower limits to thermal habitat use in this watershed). Models were fit as multi-session models pooling seasonal data from 2019, 2020, and 2021 with annually defined spatial covariates varying by session, but assuming common effects of *Temp* and *Temp + Temp^2^* across sessions. No sessional models were fit to 2018 data as both the acoustic receiver array and tagging program were in their first-year deployment phases and detections were considered preliminary as the array coverage was incomplete and the sample size was low. Model selection was conducted using Akaike’s Information Criterion with small sample correction (AICc; Burnham and Anderson 2002; Table 3). The selected model was used to predict Arctic grayling relative density of activity centres throughout the watershed as a function of the summer temperature distributions in a cool year (2019; 11.0 ± 1.1 °C) and a warm year (2021; 12.2 ± 2.0 °C). The R code for fitting and selecting models as well as extracting and visualizing activity centres are available at <*Supporting Information, Tutorial* and link>.

**Table 3.**
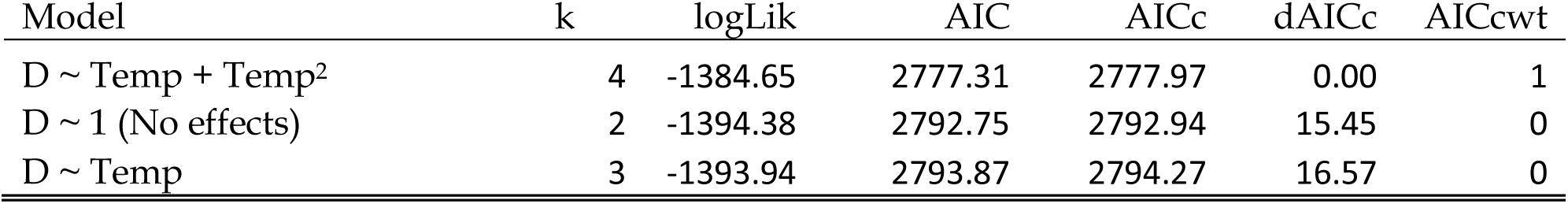
AIC statistics for candidate Arctic grayling multisession SECR models for each season. Presented are the log likelihood, the standard AIC score and the AICc score for small sample sizes, the difference in AICc between each model from the top model (ΔAICc), and the respective model weights (AICcwt). Model terms are the relative density response (D), temperature (Temp) and squared temperature (Temp^2^).

| Model | k | logLik | AIC | AICc | dAICc | AICcwt |
| --- | --- | --- | --- | --- | --- | --- |
| D ~ Temp + Temp<sup>2</sup> | 4 | -1384.65 | 2777.31 | 2777.97 | 0.00 | 1 |
| D ~ 1 (No effects) | 2 | -1394.38 | 2792.75 | 2792.94 | 15.45 | 0 |
| D ~ Temp | 3 | -1393.94 | 2793.87 | 2794.27 | 16.57 | 0 |

## Results

The acoustic receiver array recorded over three million acoustic detections that were aggregated into 5771 unique daily Arctic grayling locations for seasonal and annual daily binary observations). Model selection indicated *Temp + Temp^2^* as the best multisession model for the of relative density *D* (Table 3; *Supporting Information, Table 4*).

Relative density predictions for tagged Arctic grayling activity centres peaked at 12.4 °C (Figure 2). During the summer feeding window, tagged Arctic grayling activity centres were widely distributed throughout the Parsnip River and the four major tributaries included in this study (Figure 3). High relative densities of activity centres were estimated in the uppermost reaches of all study tributaries, and notable densities occurred in the Parsnip River mainstem near the Table River confluence.

**Figure 2.**
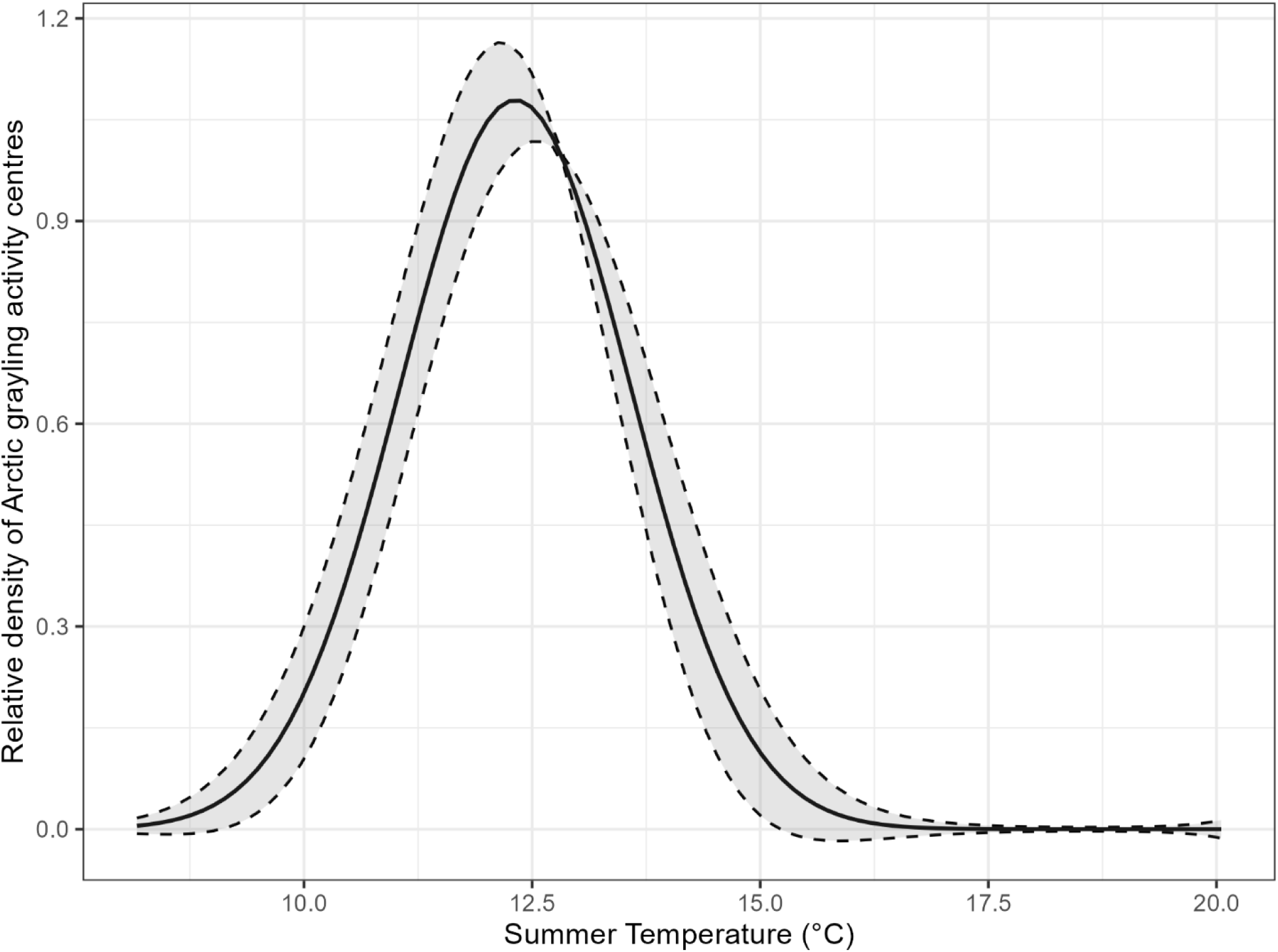
Selected model predictions of the relative density of tagged Arctic grayling activity centres with relation to the temperature covariate across all three summer seasons (panel A) Relative density of tagged Arctic grayling peaked at 12.4 °C. The shaded region indicates the± 1 standard error of model predictions.

**Figure 3.**
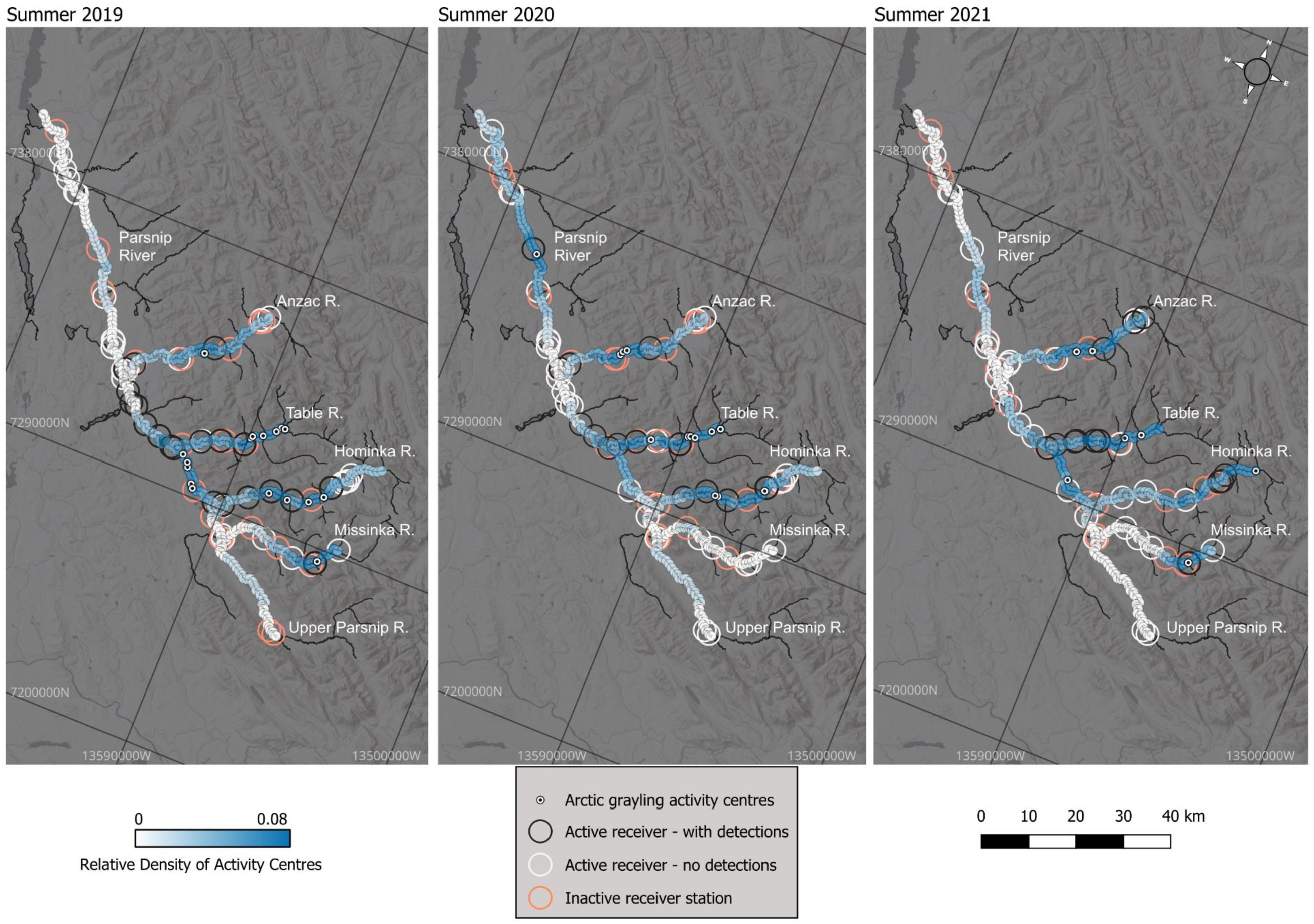
Annual summer distributions of tagged Arctic grayling activity centres estimated by the selected SECR model across the Parsnip River watershed. Activity centres are given by white markers over the relative tagged population density across the state-space in blue. Coloured circles indicate the receiver array and which acoustic receivers were active over the detection window (Jul. 1 – Sep. 15 each year). Summer activity centre distributions were wide ranging within the Parsnip River and its four main tributaries, with mainstem Parsnip River activity centres being closely associated with the Table River confluence.

Arctic grayling activity centres were widely distributed across the tributaries during the cool year (2019). However, their distribution contracted during the warm year (2021) into a patchier pattern. The relative density of Arctic grayling activity centres in the Parsnip River mainstem was predicted to be low in both years, though some river sections that were predicted to be used in the cool year were not used in the warm year.

## Discussion

The purpose of this study was to demonstrate the use of SECR models modified to accommodate relative density and tagging effort as a quantitative tool for analysing telemetry data gathered by passive automated systems. In this paper we use a case study for an aquatic species monitored using an AT detection array, though this method applies widely to both aquatic and terrestrial applications such as radio telemetry, radio frequency identification tagging, or GPS tags (e.g., Wallace et al. 2022, Shizuka et al. 2022, Taylor et al. 2017).

An attractive feature of automated telemetry data for ecologists is its ability to describe where animals distribute themselves through space and time with relative ease. A large proportion of telemetry studies to date have focused on descriptive approaches, and fewer have modeled the drivers of animal density and distribution using telemetry data (Whoriskey et al. 2019; Brownscombe et al. 2022). Formalizing an approach to telemetry analysis which quantifies relationships between spatial covariates and animal density is an important step towards unifying the vast amount of data produced by telemetry studies and the quantitative objectives that often drive resource management (Crossin et al. 2017). SECR is an effective method for achieving this end given careful consideration of how the model should be specified and interpreted when used with telemetry data.

The usual formulation of SECR models is for population estimation and we reiterate that AT data are not suited for this objective. However, here we highlighted the use of a simpler version of these models focused on the relative density as a function of spatial covariates. This formulation is appropriate for AT data and is now implemented in `secr`. We see further potential for the development of specific models for use with AT data. For example, Raabe (2013) presents a Bayesian SECR model that could be modified for use in river networks by replacing Euclidean distances with network distances. While we use a covariate that was directly measured in the field (i.e., water temperature measured with loggers and interpolated with a spatial stream network model [O’Connor et al. *in review*]), the method is useful for covariates measured using a broad range of tools (e.g., via remote sensing by satellites or drones). Further, while we address here the challenges of adapting SECR to acoustic telemetry data, our approach may be applied to any data source in which only the tagged population is available for sampling (e.g., radio telemetry, PIT tagging arrays, etc.) in both the aquatic and terrestrial environments.

We used SECR models to describe how spatial covariates influence watershed-scale distributions of the tagged Arctic grayling population. The response of tagged fish to temperature showed a peak relative density at 12.4 °C, which matched well with the results of other thermal habitat studies in this system and elsewhere. Habitat suitability indices developed in other systems for Arctic grayling suggest optimal temperature ranges of 5 – 14 °C (all life stages; Larocque et al. 2014) and 6 – 16 °C (adults; Hubert et al. 1985; Stewart et al. 2007). The sharp decline in the response at temperatures above 15 °C broadly agrees with reduced survival rates above 18 °C identified in Arctic grayling populations in the southern periphery population of Big Hole, Montana (Carillo-Longoria 2023). A study in the Parsnip River watershed found a high probability of habitat occupancy by Arctic grayling in this system at 10.9 °C (8.7 – 14.2 °C; O’Connor et al., *in review*). A series of thermal preference experiments conducted on adult Anzac River Arctic grayling found a thermal preference peak at 11.3 °C (25^th^ to 75^th^ quantile range of 8.6 – 14.1 °C; Bottoms 2024). These findings also reflect selection for cooler temperatures as found in a study of Arctic grayling habitat use by radio tagging, snorkel surveys, and thermal drone habitat mapping conducted on the Anzac River in the same year (Bottoms 2024).

When applying SECR to AT data, only tagged individuals can be detected, and it is important to consider whether the sampling design is appropriate and to verify that what is being modeled represents the distribution of animals rather than a distribution of tagging effort. The effort plug-in *P*_*T*_(***x***) represents the probability that an animal with an activity centre at ***x*** was tagged, relative to other sites. In the special case that *P*_*T*_(***x***) is uniform (i.e., representative of equal tagging probability throughout study area) it has no effect on parameter estimates. In our case study, we only had complete data on when and where tags were deployed (not actual tagging effort), and our *P*_*T*_(***x***) was a cumulative kernel smooth of these tagging locations, which inevitably included an effect of varying fish density (Figure 4). We expect this to reduce the estimated effect of covariates on density, as some of the density variation is mis-attributed to *P*_*T*_(***x***). To reduce such issues, we recommend future AT field studies to carefully record tagging effort (e.g., time spent angling at each site).

**Figure 4.**
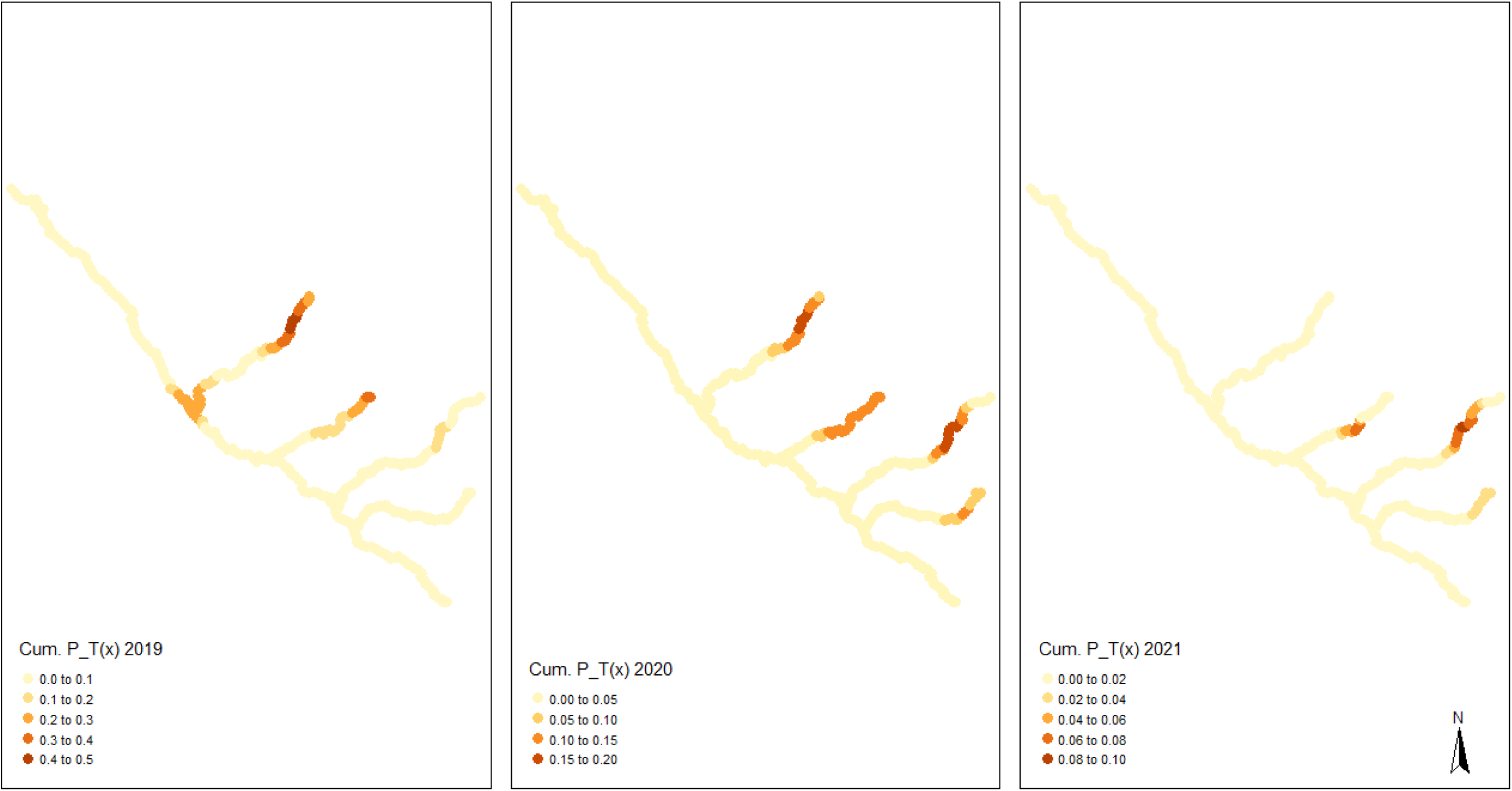
Kernel-smoothed tag deployments (*P*_*T*_(***x***)) applied by location for the summers of 2019, 2020, and 2021. Kernel density was cumulative over the first two seasons, and subtracted in the third season as tags deployed in 2018 began to reach the end of their battery life.

While we apply this approach to a dendritic river network, the underlying assumptions which apply to AT data in river networks would also apply to AT data collected in lakes, estuaries oceans, and terrestrial environments, though careful consideration should be given to how 2D state spaces differ from our case study. The assumptions of a closed population and static activity centres over each modeled window would also apply to open-water species provided that a researcher can define ecologically-relevant periods (e.g., seasons) when the individuals have settled in static home ranges. Computation times, which in our dendritic state space were relatively efficient (∼5 minutes for the most complex models on a mid-range machine), would increase in proportion to the extent of the area represented by a 2D state space, which while not prohibitive, warrants consideration among users when specifying the spatiotemporal grain and extent of their covariate pixels.

In conclusion, this work demonstrated the use of a powerful quantitative approach towards analyzing the vast amount of AT data produced by aquatic ecologists. We found that combining the statistical model with existing AT visualization techniques (e.g. animated tracks produced with the R package `RSP`; Niella et al. 2020) produced the most comprehensive understanding of Arctic grayling distribution within the available spatiotemporal dimensions of the dataset. The development of SECR models is an active field of research and has resulted in models that account for changes in the location of activity centres (Royle et al. 2014) and methods for accounting for spatial heterogeneity in detections (McLellan et al. 2023). Applying similar AT- focused modifications to these models would allow ecologists to answer a broad range of previously difficult-to-answer questions.

## Supporting information

Supplemental tutorial appendix

## Acknowledgements

We thank Murray Efford for his thoughtful inputs on this manuscript, including his help in the description of the statistical model and refinements to the secr package that were applied in our analysis. This research was funded by the Fish and Wildlife Compensation Program (grants PEA-F19-F-2593, PEA-F20-F-2961, PEA-F21-F-3178, PEA-F22-F-3388, PEA-F23-F-3652). MAM was supported by the Canadian Research Chairs program (Statistical Ecology), the BC Knowledge Development Fund, and the Canada Foundation for Innovation (John R. Evans Leaders Fund, grant 37715). MAM, EGM, MP and SJC were supported by NSERC Discovery Grants. All Arctic grayling were captured as authorized under permits PG18-356580, PG19-523435, PG20-606121, and PG21-622265 issued by the BC Ministry of Forests, Lands, Natural Resources Operations and Rural Development (currently BC Ministry of Water, Land, and Resource Stewardship). Fish handling and tagging were conducted in accordance with the University of Northern British Columbia’s Animal Care and Use Committee (ACUC protocol 2018-06).

