## Supplemental Figure 5 for "Linking acoustic telemetry data to spatial covariates in river networks with spatially explicit capture-recapture models"

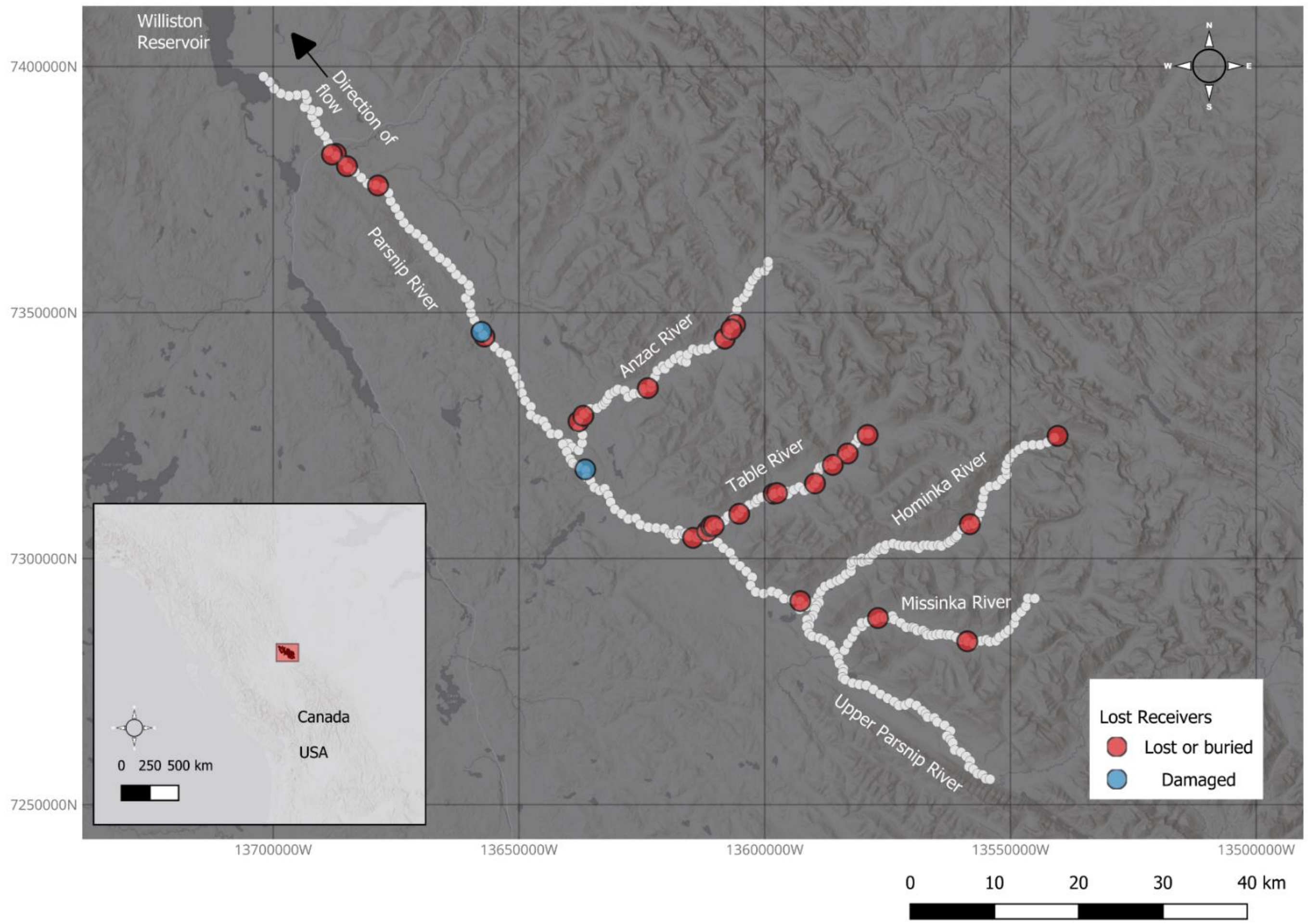


Supporting Information, Figure 5. Acoustic receivers which were lost or damaged due to high water flows. Particularly notable losses occurred in the Anzac and Table Rivers during the high discharge freshet and sustained high flows in the 2019 and 2020 seasons. Acoustic receivers were replaced at suitable sites when water levels returned to working condition
