## Supplemental Table 4 for "Linking acoustic telemetry data to spatial covariates in river networks with spatially explicit capture-recapture models"

Supporting Information, Table 4. Model coefficients for the top Arctic grayling model. Covariates Temp and $Temp^2$ represent the polynomial temperature coefficients, $\alpha$ is the detection probability, and $\sigma$ is the scale parameter. The standard error and upper and lower confidence intervals are given as SE, ucl, and lcl, respectively. The intercept is not presented here as it has no meaning in the relative density model.

| | D ~ Temp + $Temp^2$ | | | |
|---|---|---|---|---|
| Model parameters | Estimate | SE | lcl | ucl |
| Temp | -0.56 | 0.31 | -1.16 | 0.04 |
| $Temp^2$ | -1.04 | 0.28 | -1.58 | -0.49 |
| $\alpha$ | -2.49 | 0.06 | -2.60 | -2.38 |
| $\sigma$ | 9.06 | 0.02 | 9.01 | 9.10 |

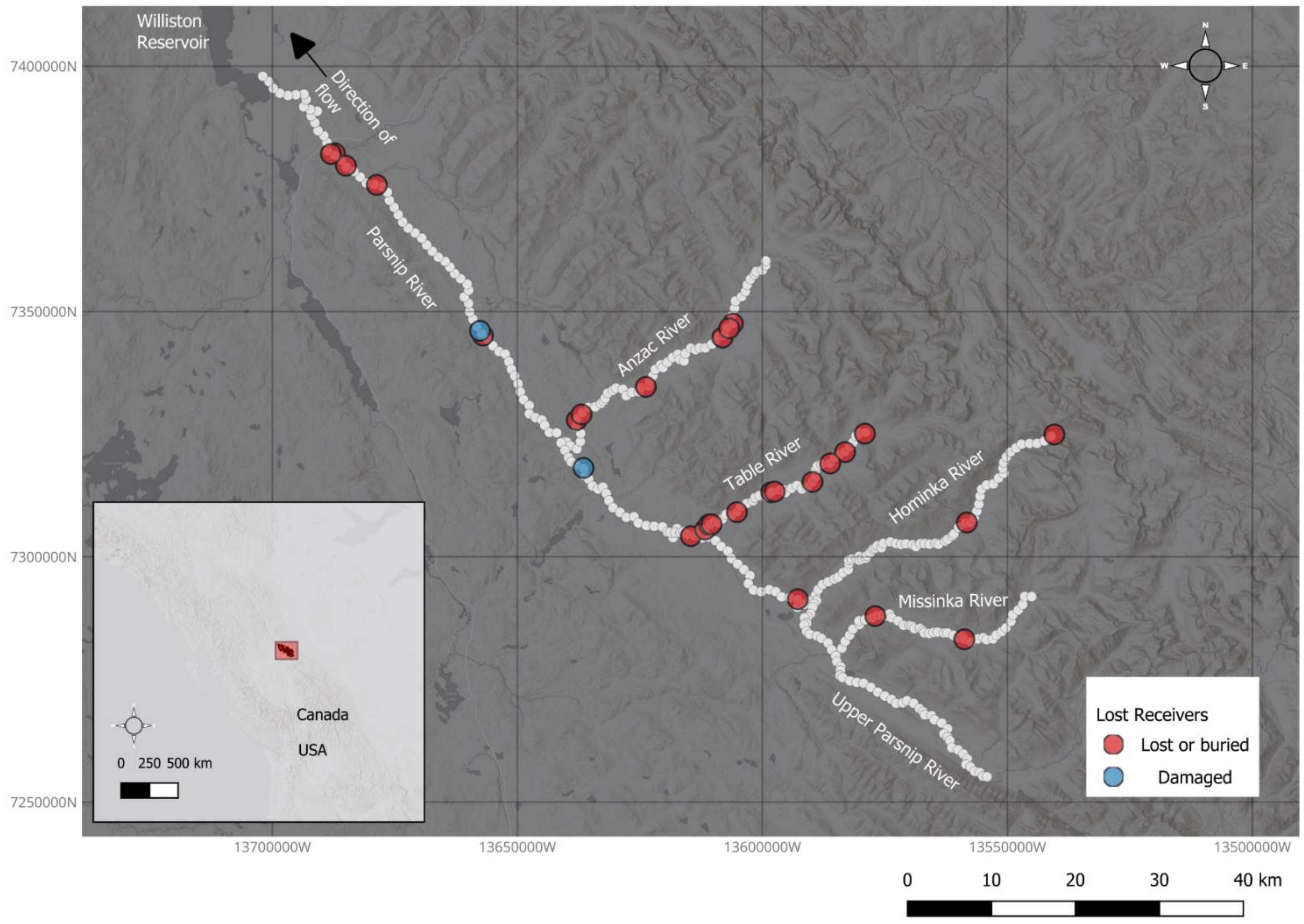


SI 2. Acoustic receivers which were lost or damaged due to high water flows. Particularly notable losses occurred in the Anzac and Table Rivers during the high discharge freshet and sustained high flows in the 2019 and 2020 seasons. Acoustic receivers were replaced at suitable sites when water levels returned to working conditions.
