## Supplemental tutorial appendix for "Linking acoustic telemetry data to spatial covariates in river networks with spatially explicit capture-recapture models"

2025-01-31

Word doc knit to submit with Supporting Information: Document is formatted properly to be hosted as a gitbook at the DOI of Archived data: <LINK WHEN LIVE>

### Introduction

The purpose of this document is to demonstrate how to link acoustic telemetry data to spatially-explicit covariates in a branching (dendritic) river network using spatially-explicit capture-recapture (SECR) models. This tutorial uses preloaded data objects, inspects the data structures necessary for the analysis, and uses the `secr` and `secrlinear` packages (Efford 2023a,b) to fit, select, predict, and visualize the models.

Our case study uses acoustic telemetry detections of Arctic grayling (*Thymallus arcticus*) from 2019-2021 in the Parsnip watershed in north-central British Columbia, Canada.

The `secrlinear` package (Efford 2023b) is a wrapper for the parent `secr` package (Efford 2023a) that enables SECR models to be fit in linear state-spaces such as dendritic river networks. Applying this approach to a 2D open water system would be similar, though care should be taken to understand how data preparation, model specifications, and computation times differ in 2D systems. Please refer to the vignettes for the `secr` package (Efford 2023a) for more.

This workflow produces a prediction of how the temperature covariate influences the summer distribution of Arctic grayling in the Parsnip watershed. While not explicitly a model of resource selection, it is most intuitively understood as a prediction of how individuals use the temperature resource within their summer home range. This model assumes demographic closure, which implies that all tags detected in a given window remain alive at the end of the window and does not account for inseason mortality or tag loss. It further assumes that there is only one activity centre for each tag during a defined ecologically-relevant period; fine-scale movements or movement parameters between seasons are not modeled by this approach. When applying this workflow, we find pre-analysis using visualization techniques provided in the `actel` (Flávio and Baktoft 2021) and RSP (Niella et al. 2020) packages helpful to ensure that the windows being modeled are ecologically-relevant to the fine-scale distributions of the focal species and meet these assumptions.

For applications using acoustic telemetry data, we found SCR to be advantageous as it (a) uses the full detection history of the tagged population to link activity centres of the tagged animals to spatially-explicit environmental covariates (analogous to second-order habitat selection in the terminology of Johnson 1980), (b) can accommodate discontinuous acoustic receiver arrays, which is beneficial in systems where acoustic receiver loss and/or redeployment between study years is characteristic, and (c) pairs well with existing descriptive approaches to acoustic telemetry analysis to offer a statistically robust tool using *a priori* information derived from these methods about a species' movement patterns.

Due to the nature of acoustic telemetry datasets in which only the tagged population is available for detection, we highlight several specific considerations and assumptions for using this approach: i.e. (a) activity centres are fixed within modeling windows by the underlying closed population model, so this method is best applied to discrete, ecologically-relevant windows in which fine-scale movements are not the focus, (b) inferences from acoustic telemetry data depict relative (not absolute) densities of only the tagged population, and (c) spatial tagging effort must be defined in the model to ensure that predictions are not merely an artifact of tagging effort across space and time.

```
# R version 4.2.3

library(secrlinear)
library(secr) # Version 5.0.0 from GitHub MurrayEfford/secr
# devtools::install_github("MurrayEfford/secr")
library(tidyverse)
library(beepr)
library(ggpubr)
library(sf)
library(kableExtra)
```

### Read and inspect prepared data files

#### Read `traps` files.

Separate `traps` objects are made for each year as acoustic receiver sites have varying usage between years. Refer to `?secr::read.traps()` for the arguments required to create a `traps` objects. These objects were prepared using the arguments `detector = 'proximity'` and `binary.usage = TRUE`.

```
traps.su19 <- readRDS('data/traps.su19.rds')
traps.su20 <- readRDS('data/traps.su20.rds')
traps.su21 <- readRDS('data/traps.su21.rds')
```

Inspect the data structure of a `traps` object's usage. Each trap (numbered row) is either active (1) or inactive (0) on each day/occasion (numbered column). In our case study, there were 76 unique acoustic receivers used across all years, but only a portion of these were active during each modeling window. The binary matrix which corresponds to a modeling session can be assigned to a `traps` object using `secr::usage(traps) <- usage.matrix`. Occasions should be indexed from 1:(number of occasions in the modeling window) for each `traps` object and not defined using a cumulative index of dates across sessions. In the example below, acoustic receiver (row) 4 was inactive until occasion (column) 11 of the 2021 summer modeling window (at which point it was deployed as a replacement for a lost receiver). By coincidence, our binary matrix was 76 rows and 76 columns, corresponding to 76 acoustic receivers and 76 days in the summer modeling window; there is no requirement for this to be a perfectly square matrix.

```
head(usage(traps.su21), 4)
```

```
##   1 2 3 4 5 6 7 8 9 10 11 12 13 14 15 16 17 18 19 20 21 22 23 24 25 26 27
28 29
## 1 1 1 1 1 1 1 1 1 1  1  1  1  1  1  1  1  1  1  1  1  1  1  1  1  1  1  1
1  1
## 2 1 1 1 1 1 1 1 1 1  1  1  1  1  1  1  1  1  1  1  1  1  1  1  1  1  1  1
1  1
## 3 0 0 0 0 0 0 0 0 0  0  0  0  0  0  0  0  0  0  0  0  0  0  0  0  0  0  0
0  0
## 4 0 0 0 0 0 0 0 0 0  0  1  1  1  1  1  1  1  1  1  1  1  1  1  1  1  1  1
1  1
##   30 31 32 33 34 35 36 37 38 39 40 41 42 43 44 45 46 47 48 49 50 51 52 53
54 55
## 1  1  1  1  1  1  1  1  1  1  1  1  1  1  1  1  1  1  1  1  1  1  1  1  1
1  1
## 2  1  1  1  1  1  1  1  1  1  1  1  1  1  1  1  1  1  1  1  1  1  1  1  1
1  1
## 3  0  0  0  0  0  0  0  0  0  0  0  0  0  0  0  0  0  0  0  0  0  0  0  0
0  0
## 4  1  1  1  1  1  1  1  1  1  1  1  1  1  1  1  1  1  1  1  1  1  1  1  1
1  1
##   56 57 58 59 60 61 62 63 64 65 66 67 68 69 70 71 72 73 74 75 76
## 1  1  1  1  1  1  1  1  1  1  1  1  1  1  1  1  1  1  1  1  1  1
## 2  1  1  1  1  1  1  1  1  1  1  1  1  1  1  1  1  1  1  1  1  1
## 3  0  0  0  0  0  0  0  0  0  0  0  0  0  0  0  0  0  0  0  0  0
## 4  1  1  1  1  1  1  1  1  1  1  1  1  1  1  1  1  1  1  1  1  1
```

#### Read detection data

```
det.dataGR <- readRDS('data/detections-gr-su.rds')
```

Inspect the detection data to see the data structures required.

- `session`: ID for which modeling session each detection belongs to. Session IDs must be named so that they sort in the same order as the sessions in the `traps` objects.

- `ID`: The code transmitted by the acoustic tag
- `occasion`: The numeric index column of the corresponding occasion in the `traps` object associated with the session
- `x`, `y`: UTM coordinates of each detection

```
glimpse(det.dataGR)

## Rows: 757
## Columns: 5
## $ session  <chr> "2019-07-01 to 2019-09-14", "2019-07-01 to 2019-09-14", "
2019…
## $ ID       <dbl> 24302, 24302, 24302, 24302, 24302, 24362, 24369, 24361, 2
4353…
## $ occasion <int> 1, 2, 5, 5, 5, 8, 8, 8, 8, 9, 10, 10, 11, 11, 11, 11, 11,
12,…
## $ x        <dbl> 567807.0, 567807.0, 563837.8, 559575.8, 557700.5, 579001.
3, 5…
## $ y        <dbl> 6061318, 6061318, 6058268, 6054844, 6054550, 6066841, 606
6841…
```

### Read covariate data

Covariates must be available for each pixel in the state-space. They can also be defined at the tag level or the acoustic receiver level, but those are not explored in this tutorial. Here we have annual mean summer temperature values (rows) for each of the 366 pixels in the Parsnip watershed state-space. The continuous temperature covariate was created using imputed temperature data in a spatial stream network model (O'Connor 2023).

```
covs <- readRDS('data/covariates.rds')

glimpse(covs)

## Rows: 366
## Columns: 5
## $ x       <dbl> 529904.3, 530234.2, 530674.6, 531439.9, 531836.0, 531876.1
, 53…
## $ y       <dbl> 6072795, 6072772, 6072335, 6071980, 6072850, 6073799, 6074
595,…
## $ temps19 <dbl> 11.68722, 11.68722, 11.70152, 11.66809, 11.62223, 11.57152
, 11…
## $ temps20 <dbl> 9.734608, 9.734803, 9.731571, 9.732794, 9.708983, 9.659940
, 9.…
## $ temps21 <dbl> 11.32968, 11.32968, 11.32185, 11.31386, 11.29521, 11.28682
, 15…

ggplot() +
  geom_sf(data = covs %>% st_as_sf(coords = c('x','y')), aes(color = temps20)
) +
  theme_bw()
```

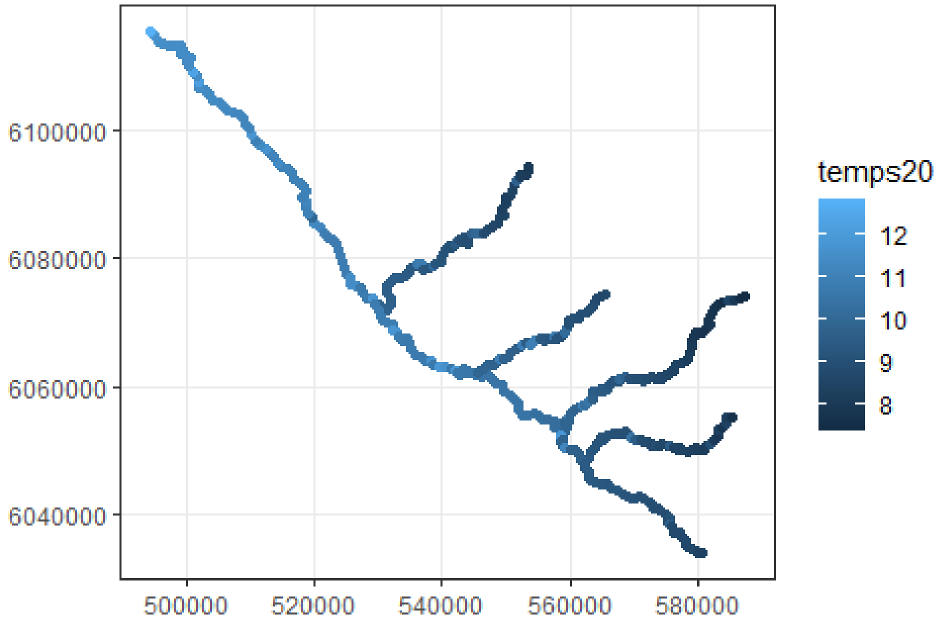


### Standardize covariates for analysis

Here we simpy standardize our covariates, as it makes model fitting go more smoothly (particularly when multiple covariates are being compared).

```
temps.all <- c(covs$temps19,
               covs$temps20,
               covs$temps21)

mtemp <- mean(temps.all)
sdtemp <- sd(temps.all)

# Standardized all years
covs <- covs %>%
  mutate(ztemp19 = (temps19 - mtemp) / sdtemp,
         ztemp20 = (temps20 - mtemp) / sdtemp,
         ztemp21 = (temps21 - mtemp) / sdtemp,
         ztemp2.19 = ztemp19^2,
         ztemp2.20 = ztemp20^2,
         ztemp2.21 = ztemp21^2)
```

### Create masks

Masks are defined for each year, as each will have unique covariate values.

```r
habitatmap <- ("data/pcr.shp") # Multiline shapefile of the Parsnip Core Regi
on (pcr)
spacing <- 1000 # Pixel size of 1,000 m

mask2019 <- read.linearmask(file = habitatmap, spacing = spacing)
mask2020 <- read.linearmask(file = habitatmap, spacing = spacing)
mask2021 <- read.linearmask(file = habitatmap, spacing = spacing)
```

Covariate columns can be assigned to `mask` objects using `covariates(mask)[,'cov.name'] <- cov.name`. Since the multi-session model fits pooled data across all sessions while assuming common covariate effects, the covariate names must be the same in each sessional mask. Also refer to `secr::addCovariates()`.

```r
covariates(mask2019)[,'ztemp'] <- covs$ztemp19
covariates(mask2020)[,'ztemp'] <- covs$ztemp20
covariates(mask2021)[,'ztemp'] <- covs$ztemp21

covariates(mask2019)[,'ztemp2'] <- covs$ztemp2.19
covariates(mask2020)[,'ztemp2'] <- covs$ztemp2.20
covariates(mask2021)[,'ztemp2'] <- covs$ztemp2.21
```

### Sampling effort

The distribution of tagging effort can bias model predictions if not accounted for. While a uniform tagging program should be applied to evenly distribute tags within the acoustic receiver array, logistical realities of sampling dynamic aquatic species and systems can at times lead to an uneven distribution of tags.

To provide the model a way to account for tagging effort in space and time and to check that any relationships found between the tags and their spatial covariates are not simply an artifact of tagging effort, a spatial effort layer is created. This layer is then added into the model as a covariate which is used to thin the likelihood (Borchers and Efford 2008) by the probability that a tag detected at a given site had been previously tagged there.

The probability surfaces used in this example were created by kernel density estimation using `qgis_run_algorithm('qgis:heatmapkerneldensityestimation')` from the `qgisprocess` package (Dunnington et al. 2023) with the kernel radius set to $\sigma$ and the weight field defined as the cumulative tags applied at each tagging pixel divided by the total tags applied to-date (inclusive of the window being modeled). From this, we subtracted the tagging efforts greater than two years old (an approximation of the ~2.2 battery life of the tags). A better metric for this object would be a cumulative time spent sampling against success rates at each pixel in the state-space.

This example accumulates effort seasonally, which in the case of Arctic grayling which show high fidelity to their summer (tagging season) habitats was reasonable. Applications of this approach should consider the ecology of a species of interest when defining an effort layer and modeling windows.

Here we visualize the summer effort layers used in each season.

```
effort <- readRDS('data/effort.rds')

ggplot() + geom_sf(data = effort, aes(color = gr.su19)) +
  theme_bw() +
  coord_sf(datum = st_crs(32610))
```

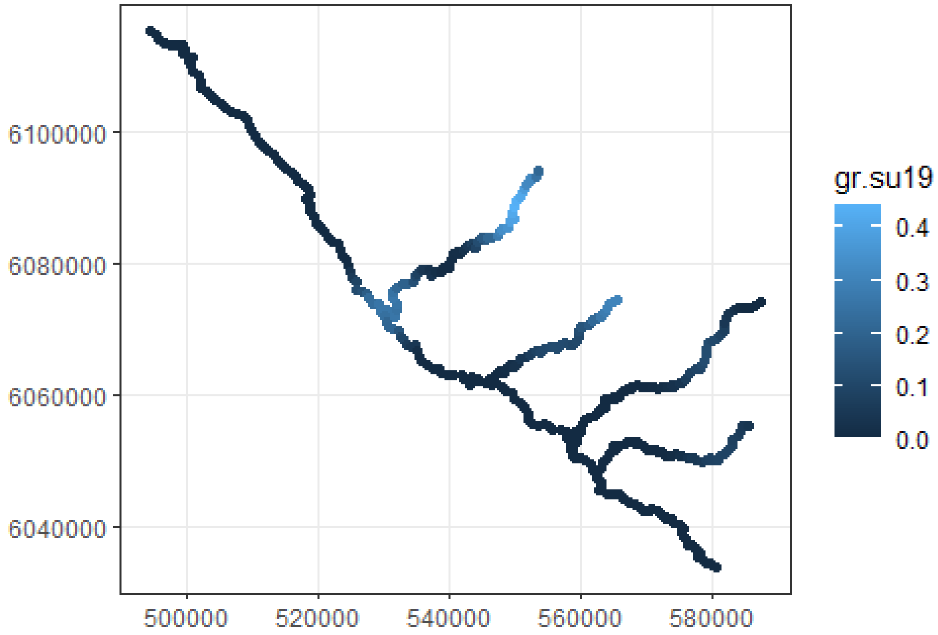


```
ggplot() + geom_sf(data = effort, aes(color = gr.su20)) +
  theme_bw() +
  coord_sf(datum = st_crs(32610))
```

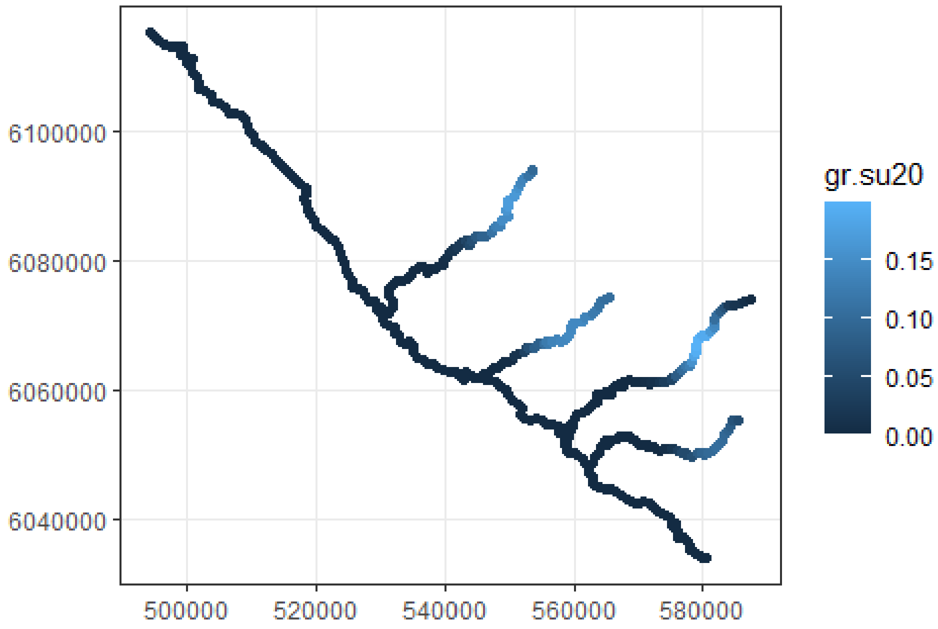


```r
ggplot() + geom_sf(data = effort, aes(color = gr.su21)) +
  theme_bw() +
  coord_sf(datum = st_crs(32610))
```

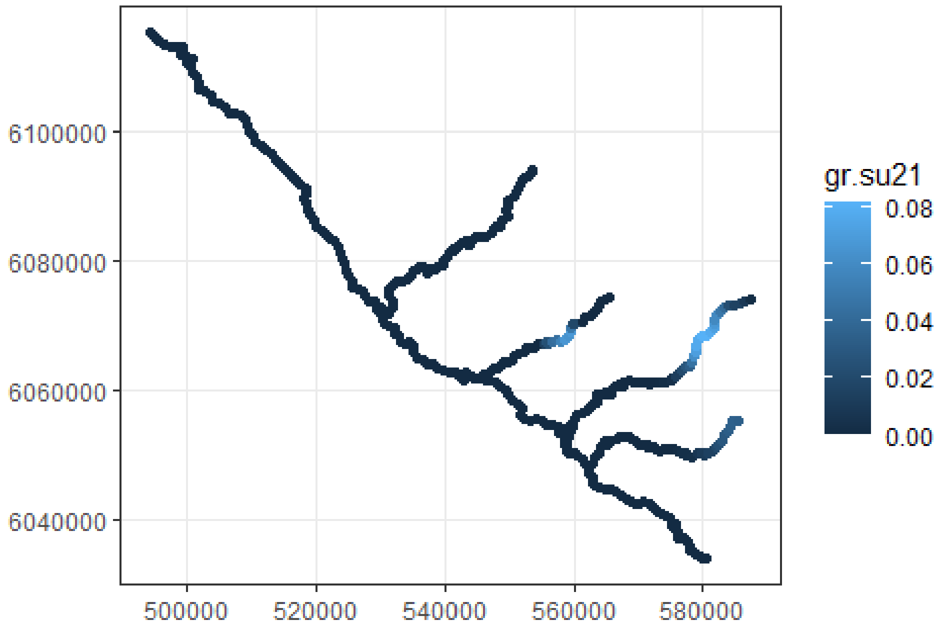

Assign effort covariates (pd for pdot in Efford's terminology) to each mask

```
covariates(mask2019)$pd <- effort$gr.su19
covariates(mask2020)$pd <- effort$gr.su20
covariates(mask2021)$pd <- effort$gr.su21
```

### Define capture history

Define the detection history of Arctic grayling as a multisession `capthist` object. Use `secr::verify` to check the integrity of the data or highlight data which is in conflict.

```
GR.su <- make.capthist(captures = det.dataGR,
                       traps = list(traps.su19,
                                    traps.su20,
                                    traps.su21),
                       fmt = 'XY',
                       bysession = TRUE)
verify(GR.su)

## No errors found :-)
```

### Fit candidate SECR models to data

Once all the data is loaded and prepared, SECR models can be fit using the arguments as defined by `?secr::secr.fit()`. Model `su.m0` is the null model, modeling `D` against the only tagging effort layer `pd`. Models `su.m1` and `su.m2` add terms for `temp` and its squared term `temp2`, respectively.

Starting parameters: `D` is 1; `g0` and `sigma` were defined after exploratory SECR modeling of acoustic telemetry detections using intercept-only models to inform appropriate starting values.

Details include the use of `networkdistance`, which enables likelihood estimation in our dendritic river network and `relativeD = TRUE`, which is required when fitting models to acoustic telemetry data in which only the relative density of tagged animals can be estimated.

The argument `steptol` is a tuning parameter for fitting the model and is not necessary in all cases. Call `?nlm()` to view the maximization error codes produced by `secr.fit()` to get a sense of which tuning parameters to define if you run into convergence issues.

```
ms.mask <- list(mask2019, mask2020, mask2021)
class(ms.mask) <- c("mask", "list")

su.m0 <- secr.fit(GR.su, mask = ms.mask,
                  model = D ~ 1,
                  trace = FALSE,
                  start = list(D = 1, g0 = 0.1, sigma = 9000),
```

```
                  details = list(userdist = networkdistance,
                                 relativeD = TRUE,
                                 externalpdot = 'pd'),
                  steptol = 1e-4)

su.m1 <- secr.fit(GR.su, mask = ms.mask,
                  model = D ~ ztemp,
                  trace = FALSE,
                  start = list(D = 1, g0 = 0.1, sigma = 9000),
                  details = list(userdist = networkdistance,
                                 relativeD = TRUE,
                                 externalpdot = 'pd'),
                  steptol = 1e-4)

su.m2 <- secr.fit(GR.su, mask = ms.mask,
                  model = D ~ ztemp + ztemp2,
                  trace = FALSE,
                  start = list(D = 1, g0 = 0.1, sigma = 9000),
                  details = list(userdist = networkdistance,
                                 relativeD = TRUE,
                                 externalpdot = 'pd'),
                  steptol = 1e-4)
```

### AICc selection of the best model

```
kable(AIC(su.m0, su.m1, su.m2) %>%
  arrange(AICc))
```

| | model | detectfn | npar | logLik | AIC | AICc | dAIC | AICwt |
|---|---|---|---|---|---|---|---|---|
| su.m2 | D~ztemp + ztemp2 g0~1 sigma~1 | halfnormal | 4 | -1384.65 | 2777.305 | 2777.972 | 0 | 1 |
| su.m0 | D~1 g0~1 sigma~1 | halfnormal | 2 | -1394.38 | 2792.751 | 2792.944 | 15.446 | 0 |
| su.m1 | D~ztemp g0~1 sigma~1 | halfnormal | 3 | -1393.94 | 2793.873 | 2794.266 | 16.568 | 0 |

### Predict the temperature covariate from the top model

Create a dataframe of new data over which to predict the model

```
covs19 <- covariates(mask2019)
covs20 <- covariates(mask2020)
covs21 <- covariates(mask2021)

npred <- 100
newdat <- data.frame(ztemp = seq(min(c(covs19$ztemp, covs20$ztemp, covs21$zte
mp)),
                                max(c(covs19$ztemp, covs20$ztemp, covs21$ztem
p)),
                                length.out = npred)) %>%
              mutate(ztemp2 = ztemp^2)
```

Predict the temperature covariate from the best model `su.m2`:

```
cov.predict.temp1 <- predict(su.m2, newdata = newdat,
                             type = "link", se.fit = TRUE, realnames = "D")

# Get predictions out of returned list object
preds.temp1 <- NULL
for(i in 1:npred){
  preds.temp1 <- rbind(preds.temp1, cov.predict.temp1[[i]])
}
```

As the data were standardized for analysis, here we provide the non-transformed temperature variable from `covs` so that the x-axis of the plot is sensible.

```
predictions.temp <- bind_cols(preds.temp1, newdat) %>%
  mutate(temp = sdtemp * ztemp + mtemp,
         D_exp = exp(estimate),
         D_se_lwr = exp(estimate - SE.estimate),
         D_se_upr = exp(estimate + SE.estimate),
         D_95ci_lwr = exp(estimate - 1.96 * SE.estimate),
         D_95ci_upr = exp(estimate + 1.96 * SE.estimate))
```

Plot the response in Arctic grayling relative tag density to temperature:

```
covplot.temp2.su.se <- ggplot(predictions.temp, aes(x = temp, y = D_exp)) +
  geom_line(linewidth = .7) +
  geom_line(aes(x = temp, y = D_se_upr), linetype = 2) +
  geom_line(aes(x = temp, y = D_se_lwr), linetype = 2) +
  geom_ribbon(aes(x = temp, ymin = D_se_lwr, ymax = D_se_upr),
              fill = "gray50", alpha = 0.2) +
  labs(x = "Summer Temperature (°C)", y = "Relative density of Arctic graylin
g activity centres") +
  theme_bw()

covplot.temp2.su.se
```

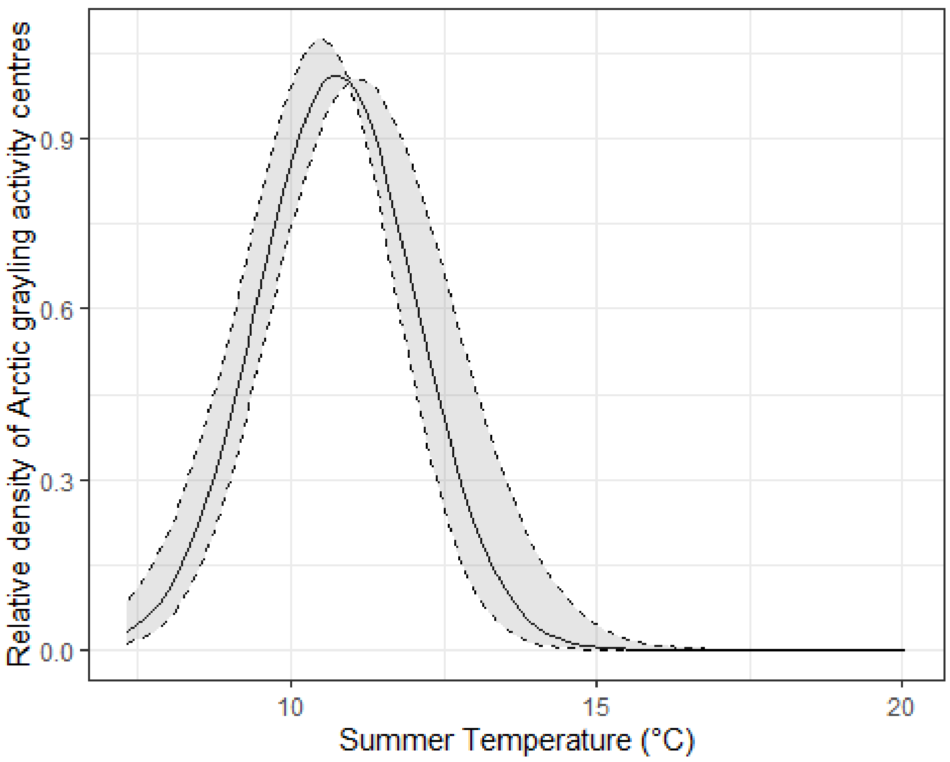


### Plotting model spatial outputs

This section plots model predictions of the relative densities and activity centres of tagged Arctic grayling during each summer season.

#### Data prep

Get session-specific model outputs:

```
m2.19 <- update(su.m2, capthist = GR.su[[1]], mask = mask2019)
m2.20 <- update(su.m2, capthist = GR.su[[2]], mask = mask2020)
m2.21 <- update(su.m2, capthist = GR.su[[3]], mask = mask2021)
```

Here we represent activity centres for each tag as the pixel with the maximum probability of detection.

```
centers19 <- fxi(m2.19)

## using external pdot

tags19 <- names(centers19)
act.cent19 <- data.frame(tag = 1:length(centers19),
                         x = NA,
                         y = NA)
for (i in 1:length(centers19)) {
  tmp2 <- max(centers19[[i]]) # Find maximum value in pdf for tag i
```

```
  tmp3 <- match(tmp2, centers19[[i]]) # Match value to index
  tmp4 <- mask2019[tmp3,] # Assign x-y coordinates of activity centre
  act.cent19[i,]$tag <- tags19[i] # Populate dataframe rows
  act.cent19[i,]$x <- tmp4$x
  act.cent19[i,]$y <- tmp4$y
}

centers20 <- fxi(m2.20)

## using external pdot

tags20 <- names(centers20)
act.cent20 <- data.frame(tag = 1:length(centers20),
                        x = NA,
                        y = NA)
for (i in 1:length(centers20)) {
  tmp2 <- max(centers20[[i]]) # Find maximum value in pdf for tag i
  tmp3 <- match(tmp2, centers20[[i]]) # Match value to index
  tmp4 <- mask2020[tmp3,] # Assign x-y coordinates of activity centre
  act.cent20[i,]$tag <- tags20[i] # Populate dataframe rows
  act.cent20[i,]$x <- tmp4$x
  act.cent20[i,]$y <- tmp4$y
}

centers21 <- fxi(m2.21)

## using external pdot

tags21 <- names(centers21)
act.cent21 <- data.frame(tag = 1:length(centers21),
                        x = NA,
                        y = NA)
for (i in 1:length(centers21)) {
  tmp2 <- max(centers21[[i]]) # Find maximum value in pdf for tag i
  tmp3 <- match(tmp2, centers21[[i]]) # Match value to index
  tmp4 <- mask2021[tmp3,] # Assign x-y coordinates of activity centre
  act.cent21[i,]$tag <- tags21[i] # Populate dataframe rows
  act.cent21[i,]$x <- tmp4$x
  act.cent21[i,]$y <- tmp4$y
}
```

### Create plots

Working with relative densities in acoustic telemetry with multiple sessions may require some fine tuning of the color palette relative to the densities predicted in each session. The built-in plotting functions are useful for technical work and the spatial data can be saved as shapefiles for making further customized visualizations in a preferred GIS using the package `sf` (Pebesma and Bivand 2023, Pebesma 2018).

```
cols19 <- terrain.colors(n = 13, rev = TRUE)
plot(predictDsurface(m2.19), col = cols19, title = "Relative D")
```

```
## Warning: too few colours; using terrain.colors(15)

points(act.cent19$x, act.cent19$y)
```

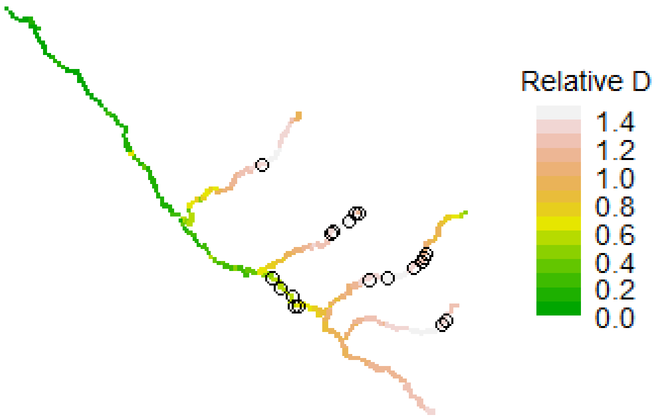


```
cols20 <- terrain.colors(n = 7, rev = TRUE)
plot(predictDsurface(m2.20), col = cols20, title = "Relative D")

## Warning: too few colours; using terrain.colors(11)

points(act.cent20$x, act.cent20$y)
```

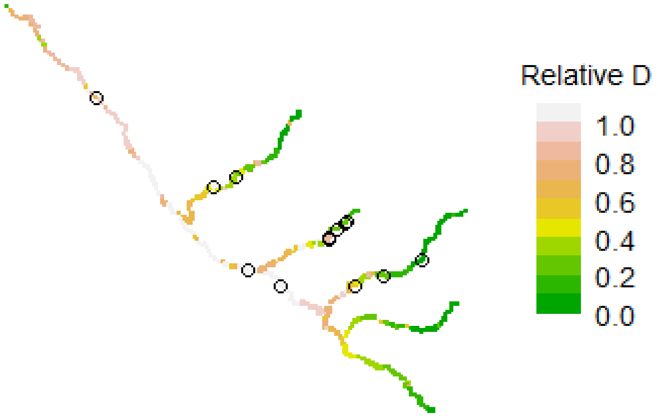


```
cols21 <- terrain.colors(n = 11, rev = TRUE)
plot(predictDsurface(m2.21), col = cols21, title = "Relative D")

## Warning: too few colours; using terrain.colors(14)

points(act.cent21$x, act.cent21$y)
```

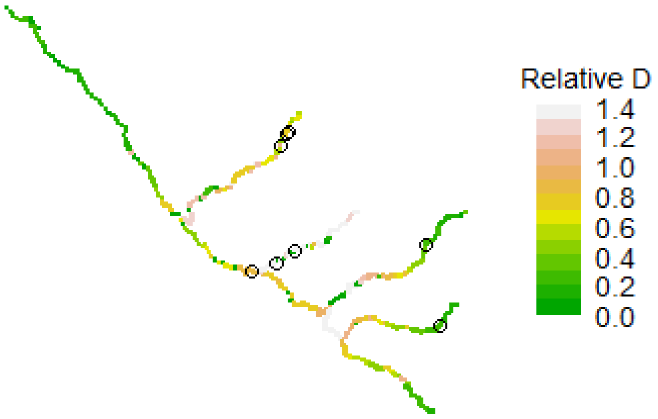


### Save outputs for visualization in GIS

Ensure that the GIS project is closed (or shapefiles are unloaded in the GIS environment) when running this code so that files can update.

```
tag.cen19 <- act.cent19%>%
  st_as_sf(coords = c('x', 'y'), crs = 32610)
rd.19 <- predictDsurface(m2.19)
tag.den19 <- rd.19 %>%
  st_as_sf(coords = c('x', 'y'), crs = 32610) %>%
  mutate(D.0 = covariates(rd.19)$D.0)

write_sf(tag.den19, 'outputs/GRtag.den2019.shp')
write_sf(tag.cen19, 'outputs/GRtag.cen2019.shp')


tag.cen20 <- act.cent20%>%
  st_as_sf(coords = c('x', 'y'), crs = 32610)
rd.20 <- predictDsurface(m2.20)
tag.den20 <- rd.20 %>%
  st_as_sf(coords = c('x', 'y'), crs = 32610) %>%
  mutate(D.0 = covariates(rd.20)$D.0)

write_sf(tag.den20, 'outputs/GRtag.den2020.shp')
write_sf(tag.cen20, 'outputs/GRtag.cen2020.shp')
```

```
tag.cen21 <- act.cent21%>%
  st_as_sf(coords = c('x', 'y'), crs = 32610)
rd.21 <- predictDsurface(m2.21)
tag.den21 <- rd.21 %>%
  st_as_sf(coords = c('x', 'y'), crs = 32610) %>%
  mutate(D.0 = covariates(rd.21)$D.0)

write_sf(tag.den21, 'outputs/GRtag.den2021.shp')
write_sf(tag.cen21, 'outputs/GRtag.cen2021.shp')
```
